# Cell-type specialization of layer 5 excitatory neurons in tactile behavior

**DOI:** 10.1101/2024.03.15.585205

**Authors:** Samson G. King, Phillip Maire, Adam Mergenthal, Stefanie Walker, Samuel Andrew Hires

## Abstract

Layer 5 is the canonical output layer of sensory cortex. The two most numerous neural constituents of Layer 5 are pyramidal tract (PT) and intratelencephalic (IT) neurons. These output cell classes combine diverse sets of inputs and project to distinct locations across the brain, suggesting differing roles in sensory information processing. Here, we investigated the representation of touch and whisker motion in these two cell types within primary somatosensory cortex (S1) using optogenetically targeted single unit electrophysiology during whisker-guided object localization. PT neurons (N = 32) had much higher spike rates than IT (N=26) during behavior. Individual members of both were modulated by, but average population firing rates were stable between quiet and whisking periods. PT neurons showed greater absolute spike rate changes, but less relative modulation than IT neurons to whisking kinematic features. Touch-excited PT (N = 18) and IT neurons (N = 8) rapidly adapted to active touch. Both populations encoded the azimuthal position of touched objects, with IT neurons more sharply tuned to position. However, position was more precisely decodable from PT population activity, due to greater evoked spikes per touch. A consequence of these characteristics is that PT neurons, with their higher firing rates, may be more effective participants in rate-based neural codes, while IT neurons, with their sharp modulation, may be more effective in timing or synchrony-based codes.

## Introduction

The brain’s ability to locate objects in space is a fundamental function for coordinated motor behavior. Precisely how object localization is encoded at the level of the cortex is still being unraveled. For touch-guided localization in rodents, one of the potential ways the brain may do this in the primary somatosensory cortex (S1) is by integration of an efference copy of motor information arriving from a separate pathway (e.g. primary motor cortex; M1) that could disambiguate the precise position of an object touched with whiskers (Hill et al., 2011). Alternatively, all information necessary for disambiguating position could arise from direct projections from the ventral posteromedial nucleus (VPM) to laminae where touched-object location is encoded (Hires et al., 2015; Petreanu et al., 2009; Armstrong–James et al., 1992; Fee et al., 1997).

In either case, we hypothesized that the cells most likely to faithfully encode touched-object location are layer 5 (L5) excitatory neurons, particularly L5 pyramidal tract (PT) cells. Neurons of this class receive broad, diverse inputs from cortical (LeFort et al., 2009; Hooks et al., 2011) and sensory input from subcortical (Petreanu et al., 2009) sources. They possess location-dependent information represented in their distal apical tufts (Xu et al., 2012; Petreanu et al., 2012) and soma (Cheung et al., 2020; Ranganathan et al., 2018). They receive local input from several other layers in the cortex but likely contribute little to local computations in the circuit since their axons seldom branch within the cortex (Narayanan et al., 2015). PTs project to many subcortical targets (Guo et al., 2017) such as the striatum (Hooks et al., 2018; Hintiryan et al., 2016; Wall et al., 2013) and the posterior medial thalamic nucleus (POm) (Sumser et al., 2017; Groh et al., 2014). This placement in the circuit makes PT neurons uniquely well-suited to integrating sensory information required for disambiguating touched-object location.

Our prior work identified a population of L5 excitatory neurons in S1 whose activity was enhanced during object touch at preferred positions that tiled the available sample space (Cheung et al., 2020). Furthermore, the pole location could be accurately predicted (60.5% at ≤ 0.5mm distance) from these touch responses using a multinomial generalized linear model (GLM) (Cheung et al., 2020). However, our previous approach did not differentiate between the candidate cell type we hypothesized to be most likely to represent object location in S1, L5 PT neurons, and the other major output cell type in L5, intratelencephalic (IT) neurons.

Here, we used two mouse line crosses that selectively express ChR2(H134R) in L5 PTs or L5 ITs to opto-tag and disambiguate the cellular class of the collected L5 pyramidal cell activity. Following identification, we employed single-unit juxtacellular electrophysiology to investigate neural representations of whisking and touch during active behavior. We found that PTs and ITs had many units that were either positively or negatively modulated by whisking, indicating that both cell types encode self-motion. Both cell types also displayed units that responded to touch at similar latencies and for similar durations, with PTs displaying much larger absolute touch responses. At the same time, many ITs showed a greater percent increase above their on-average low firing rate. ITs and PTs exhibited touch adaptation to serial pole sampling within a given trial and had attenuated responses at smaller inter-touch intervals (ITIs). However, touch-excited ITs did have a more rapid drop-off in firing rate after the first touch. Both PTs and ITs could decode object location with a GLM, with PTs outperforming both ITs alone and a mixed cell-type decoder. These data suggest that PTs are effective at representation of object location with a rate-based neural code. On the other hand, the low-latency transient touch responses in ITs may be better suited for timing or synchrony-based codes.

## Results

### Experimental Design

To investigate potential cell-type specialization of excitatory layer 5 neurons for tactile sensation and object localization, we combined loose-seal juxtacellular electrophysiology and high-speed videography during behavior. Water-restricted mice (n = 23) were trained to perform a single whisker-guided go/no-go object localization task (Cheung et al., 2019). Mice were required to investigate the position of an actuated pole and report their percept via a lick to attain a water reward or withhold licking to avoid a time-out (**Figure 1A-C**). Whiskers were tracked at 1000 frames-per-second (fps), traced, and decomposed into constituent whisker kinematic variables (**Materials and Methods**).

**Figure 1.**
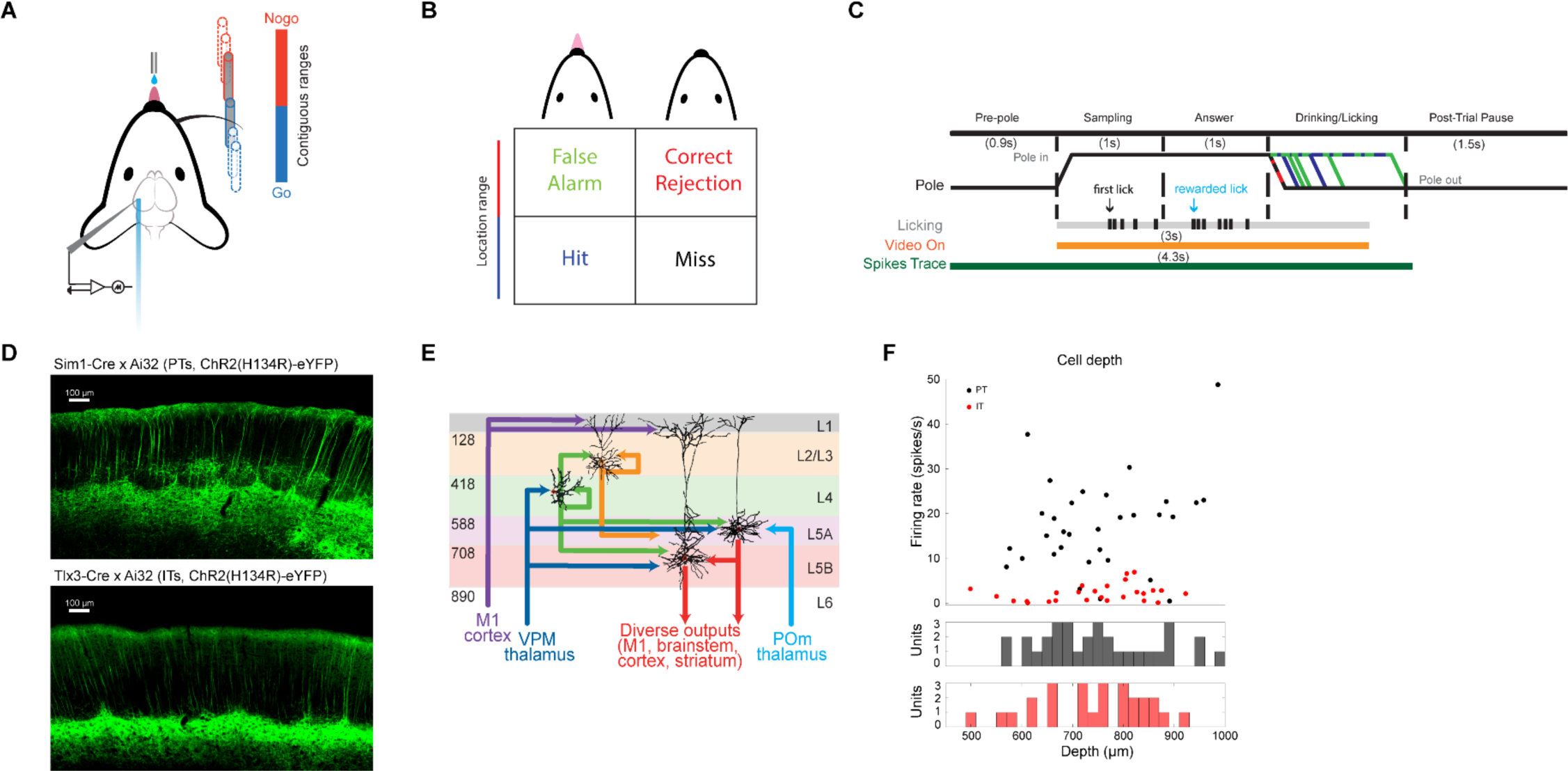
Head-fixed task, Circuitry, and Cell Depth. (A) Task schematic. To receive a reward, mice are trained to discriminate between a near (go) and far (nogo) range. Opto-tagging was performed via 10Hz, 10ms-wide pulses of 473 nm light prior to further data collection. (B) Diagram of possible trial outcomes (hit, miss, false alarm, correct rejection). (C) Trial structure and aligned data streams. Once recording had begun, high-speed (1000 fps) videography and precise electrophysiology were collected alongside behavior. The trial structure differed in ending times depending on the choice selection behavior of individual mice. (D) Histology showing expression of ChR2(H134R)-eYFP in either PT (top, Sim1-cre x Ai32 cross) or IT (bottom, Tlx3-cre x Ai32 cross) cells. (E) Circuit diagram displaying some of the input and output pathways for PT (left) and IT (right) L5 excitatory cell types in S1. Modified from Lefort et al., 2009. (F) Depths of collected cells plotted alongside their mean firing rate (top). The depth distributions of PT cells (black, mean depth 751.5 µm, range 570-986 µm) and IT cells (red, mean depth 742.2 µm, range 498-922 µm) are shown in the middle and bottom subplots, respectively.

Loose-seal (∼7.6 MOhms) juxtacellular electrophysiological recordings were performed following intrinsic signal imaging targeted to the C2 primary whisker barrel and subsequent craniotomy (**Supplemental Figure 1A,B**). All recordings were performed using one of two mouse line crosses, each of which respectively expressed channelrhodopsin-2 (ChR2-H134R) in either layer 5 PT (Sim1-cre x Ai32, n = 14 mice) neurons or layer 5 IT (Tlx3-cre x Ai32, n = 9 mice) neurons (Gerfen et al., 2013; **Figure 1C, D**). We recorded a total of 135 neurons between the two mouse lines (73 Sim1-cre x Ai32 / 62 Tlx3-cre x Ai32), which were opto-tagged using 472 nm pulsed (10Hz, 10ms width) light directed onto the site of the craniotomy (∼200 µm).

Since there is a known unidirectional projection from IT neurons to PT neurons (Kiritani et al., 2012; Groh et al., 2010; Anderson et al., 2010; Brown & Hestrin, 2009), we employed stringent post-illumination spike latency (time to first spike <= 5.6 ms) and waveform characteristics to identify types (**Methods; Supplemental Figure 2A-D**). Following this approach, we were left with a population of 32 putative PT and 26 putative IT neurons. Recordings of PT neurons spanned a range of depth from pia (570-986 µm, mean 751.5 µm), while IT recordings were somewhat shallower on average (498-922 µm, mean 742.2 µm; **Figure 1E**). Across the recording session, the PT population had significantly higher mean firing rates (population mean: 17.42 ± 3.53 spk/s) than the IT population (population mean: 2.23 ± 0.73 spk/s; p = 9.91e-11, Two-Sample Kolmogorov-Smirnov Test) (**Supplemental Figure 3**), with no discernable trend of either with estimated cell body depth. This difference in firing rate demonstrates that the greater excitability of PT over IT neurons (Hattox & Nelson, 2007; Kasper et al., 1994) and their propensity for different firing modes (Chagnac-Amitai et al., 1990) translate into higher relative firing rates for PT neurons when these cortical circuits are engaged in active tactile exploration.

### Whisking-mediated activity

Given the distinct input circuitry to PT and IT neurons in S1 (Mao et al., 2011; Constantinople & Rudy, 2013) (**Figure 1D**), we reasoned that there might be fundamental differences in their responsiveness to whisker motion when viewed independently rather than as a combined class of cells. This notion is supported by previous work in which putative ITs in L5a of rat cortex displayed enhanced firing rates compared to PTs during whisking periods with no contacts (de Kock & Sakmann, 2009). For both classes, the majority of identified neurons showed significant modulation to whisking (PT 29/32; IT 21/26 chi-squared test, 95% significance), with a balanced distribution of neurons that were positively versus negatively modulated (**Figure 2A, B**). However, for both cell types, the net change in population spiking between quiet and whisking periods was not significantly different from zero (PTs: 0.83 ± 5.41Δ spks/s p = 0.33; ITs: 0.067 ± 1.82 Δ spks/s p = 0.38, Wilcoxon signed-rank test **Figure 2A-C**). The absolute modulation expressed as changes in spike rate was significantly larger in PT than in IT neurons (**Figure 2B, C**; 3.9 spks/s for PTs vs 1.1 spks/s for ITs p = 2.51e-05, Mann-Whitney U Test). However, IT neurons were more relatively modulated than PTs (mean 0.32, median 0.28, modulation depth for ITs vs. mean 0.16, median 0.11 modulation depth for PTs p = 0.01, Mann-Whitney U Test; **Figure 2D, E**). Overall, the gross firing characteristics of PT and IT neurons between quiet and whisking periods primarily differed in the mean spike rate of the two populations rather than modulation characteristics.

**Figure 2.**
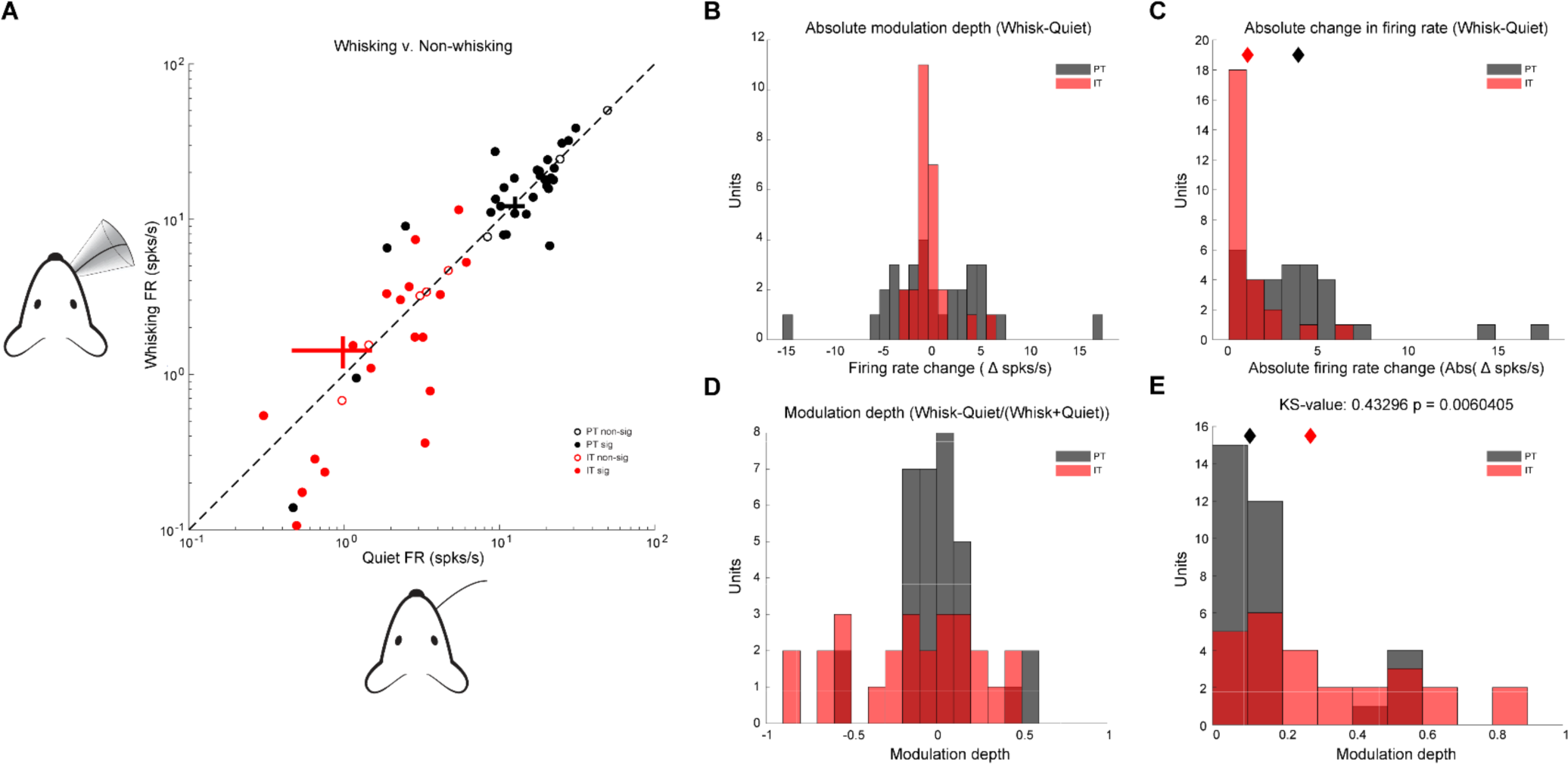
Whisking Modulation of Firing Rate. (A) Schematic, y-axis, displaying whisking. Schematic, x-axis, displaying quiescence. Firing rate for non-whisking (PTs: 16.30 ± 9.93 spks/s; ITs: 2.23 ± 1.70; mean + sd) and whisking periods (PTs, black: 17.14 ± 10.59 spks/s; ITs, red: 2.30 ± 2.67 spks/s; mean ± sd) PTs: p = 0.33, z-value = 0.97, signed rank = 316; ITs: p = 0.38, z-value = −0.88, signed rank = 141, Wilcoxon signed-rank test). Significantly whisking-modulated cells (Methods and Materials) are filled circles, non-significant cells are open. (B) Absolute modulation depth, displayed as the firing rate during quiescent periods subtracted from the firing rate during active whisking periods (PTs, black; ITs, red). (C) As in (B), except all values are displayed as absolute values. Means for each population are displayed as diamonds in their respective colors (PTs, black: 3.9 spks/s; ITs, red: 1.1 spks/s, p = 2.51e-05, z-value = 4.21, rank sum = 1214, Mann-Whitney U Test). (D) Modulation Depth (see Materials and Methods) comparing the modulation of cells between whisking and quiescent periods. PT cells’ modulation depths are displayed in black, IT cells’ modulation depths are displayed in red. Bin width for visualization: 0.1. (E) As in (D), except all values are displayed as absolute values. Means for each population are displayed as diamonds in their respective color (PTs, black: 0.16 mean/0.11 median modulation depth, ITs, red: 0.32 mean/0.28 median modulation depth, p=0.01, z-value = −2.51, rank sum = 783, Mann-Whitney U Test).

### Free-Whisking Kinematics

Despite the absence of change in population-averaged spike rates between quiet and whisking periods, the timing of spikes during whisking could be structured with respect to the whisker angle and associated kinematic variables (angle, amplitude, midpoint, and phase; **Figure 3A**). Tuning curves of single cells (**Figure 3B**) showed significant modulation across these parameters for both cell types. Firing rates were most modulated by angle (PTs: 8.81 mean Δ spks/s; ITs: mean Δ 1.63 spks/s) and midpoint (PTs: 8.93 mean Δ spks/s change; ITs: 2.01 mean Δ spks/s change), and less modulated by amplitude (PTs: 6.29 mean Δ spks/s; ITs: 1.37 mean Δ spks/s) and phase (PTs: 5.05 mean Δ spks/s; ITs: 0.8967 mean Δ spks/s) (**Figure 3C**). Overall, PT neurons had significantly greater absolute modulation (angle: p = 6.59e-08; phase: p = 5.07e-08; amplitude: p = 1.21e-06; midpoint: p = 1.56e-07, Mann-Whitney U-test; **Figure 3C**), while IT neurons showed significantly higher relative modulation to each kinematic variable (angle: p = 4.77e-03; phase: p = 2.02e-03; amplitude: p = 2.48e-03; midpoint: p = 1.32e-04, Mann-Whitney U-test; **Figure 3D, Supplemental Figure 4A-C**). The finding that midpoint provided the greatest modulation depth for both PT and IT neurons was consistent with prior results in undifferentiated L5 excitatory neurons (Cheung et al., 2020) and underscores its potential importance in whisker-mediated behaviors (Cheung et al., 2019).

**Figure 3.**
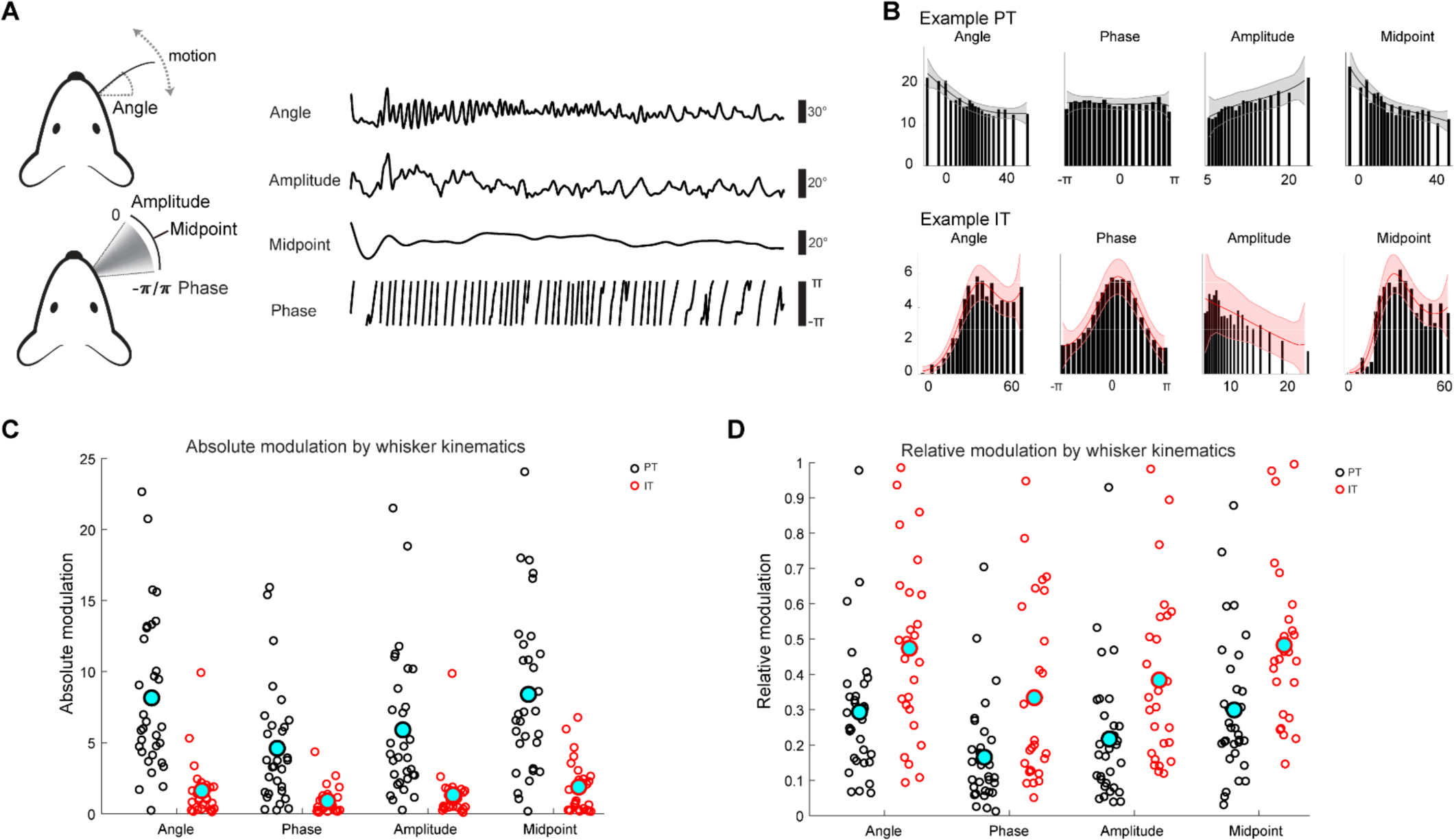
Whisker Kinematic Variable Spiking Modulation. (A) Schematic of a mouse (left) sweeping its whisker back and forth during an active whisking period with the concomitant kinematic variables displayed alongside. On the right, whisker kinematics extracted using a Hilbert transform are displayed, with scale bars relevant to each. (B) PT example unit (top) and IT example unit (bottom), each showing their respective tuning curves to angle, phase, amplitude, and midpoint. (C) Absolute modulation of PTs (black) and ITs (red) to each kinematic variable displayed as a swarmplot. Means for each group are displayed as a large circle with a cyan fill. All statistical comparisons use a Mann-Whitney U Test. Angle: PT mean = 8.15 Δ spks/s, IT mean = 1.62 Δ spks/s; p = 6.59e-08, z-value = 5.40, rank sum = 1290. Phase: PT mean = 4.60 Δ spks/s, IT mean = 0.90 Δ spks/s; p = 1.41e-06, z-value = 4.82, rank sum = 1253. Amplitude: PT mean = 5.90 Δ spks/s, IT mean = 1.32 Δ spks/s; p = 2.18e-07, z-value = 5.18, rank sum = 1276. Midpoint: PT mean = 8.40 Δ spks/s, IT mean = 1.88 Δ spks/s; p = 5.41e-07, z-value = 5.01, rank sum = 1265. (D) Relative modulation of PTs (black) and ITs (red) to each kinematic variable displayed as a swarmplot. Means for each group are displayed as a large circle with a cyan fill. All statistical comparisons use a Mann-Whitney U Test. Angle: PT mean modulation depth = 0.29, IT mean modulation depth = 0.47; p = 3.5e-03, z-value = −2.92, rank sum = 757. Phase: PT mean modulation depth = 0.17, IT mean modulation depth = 0.33; p = 5.5e-03, z-value = −2.78, rank sum = 766. Amplitude: PT mean modulation depth = 0.22, IT mean modulation depth = 0.38; p =1.7e-03, z-value = −3.13, rank sum = 743. Midpoint: PT mean modulation depth = 0.48, IT mean modulation depth = 0.30; p = 8.44e-04, z-value = −3.34, rank sum = 730.

Within individual PT and IT neurons, there was a substantial degree of variability in modulation depth across particular kinematic variables; however, ITs with lower firing rates tended to have similar modulation depths between different kinematic variables (**Figure 4A**). Across the population, PTs’ phase responses correlated the least well with other kinematic variables (angle: R = 0.67, p = 2.7e-05; amplitude: R = 0.65, p =5.4e-05; midpoint: R = 0.60, p = 2.9e-04, Pearson’s R correlation). PTs’ angle responses, on the other hand, correlated well with the non-phase kinematic variables (amplitude: R = 0.85, p = 9.1e-10; midpoint: R = 0.84, p = 2.5e-09, Pearson’s R correlation) (**Figure 4B, left**). The IT population’s phase-midpoint pair had the lowest correlation (R = 0.39, p = 0.044), closely followed by its amplitude-midpoint pair (R = 0.48, p = 0.014). Except for midpoint, ITs’ phase correlations with other variables tended to be stronger than those of PTs’, with the second strongest association between it and angle (R = .75, p = 1.2e-5). The strongest association was observed between angle and amplitude (R = 0.79, p = 1.4e-06) (**Figure 4B, right**). Angle and amplitude modulation are well-correlated in both cell classes, while phase and midpoint are more poorly correlated. The commonalities in co-variation across cell types could indicate that the pathways carrying slowly-varying (amplitude and midpoint) and quickly-varying (angle and phase) kinematic variables’ information converge on both L5 output cell classes.

**Figure 4.**
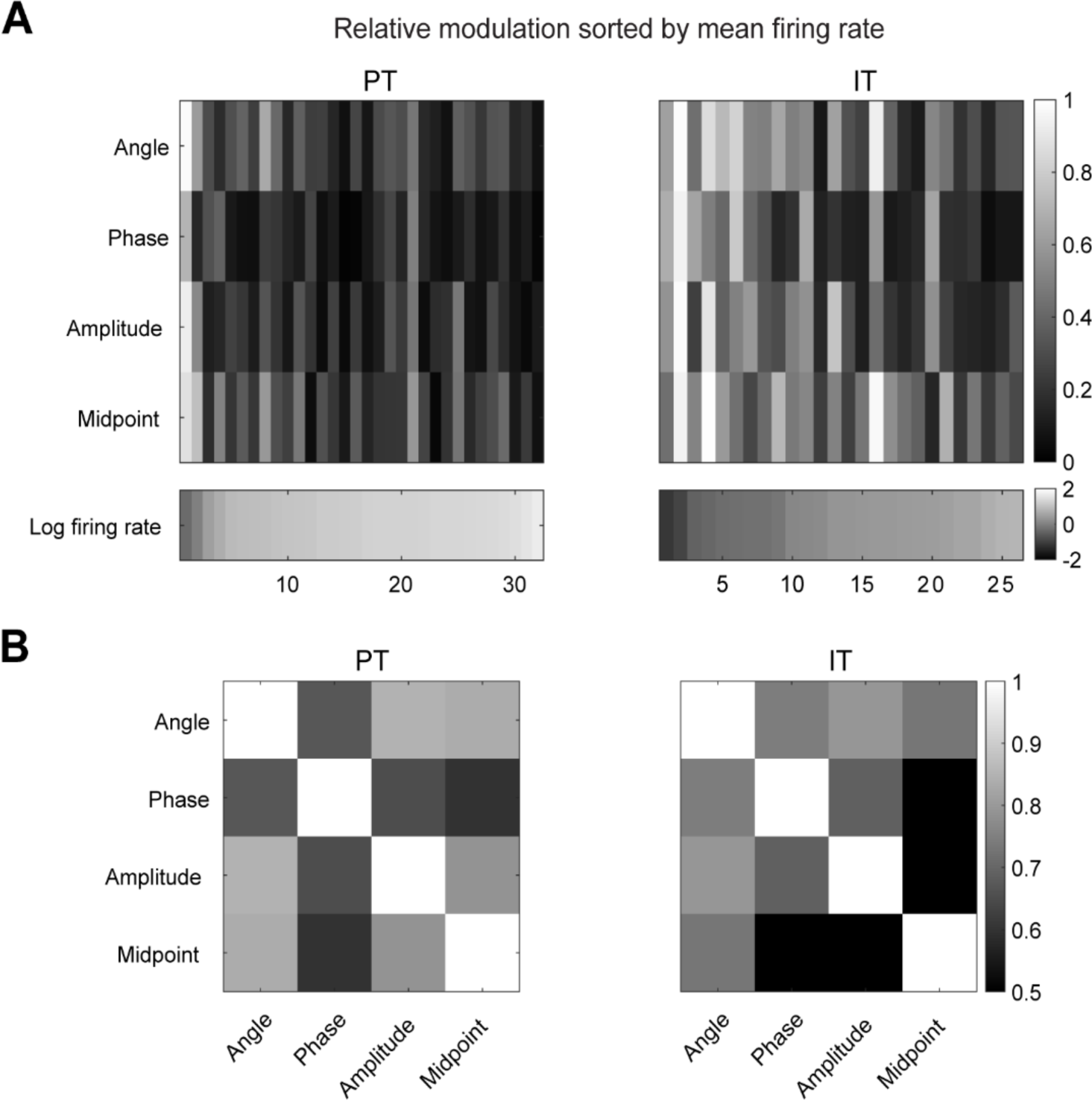
Population Modulation and Correlation Between Kinematic Variables. (A) Relative modulation of PTs (left) and ITs (right) by angle, phase, amplitude, and midpoint sorted by firing rate (displayed in log scale at the bottom for each cell type). (B) Contingency tables of Pearson correlation coefficients for modulation depths across angle, phase, amplitude, and midpoint for PTs (left) and ITs (right).

### Touch Responses

Across the recorded populations, 78% (25/32) of PT and 46% (12/26) of IT neurons were transiently modulated by touch (**Figure 5A, B**). Of the modulated population of PTs, 72% (18/25) were excited at the time of touch, and 28% (7/25) were inhibited. A similar percentage of touch-modulated ITs were excited (67%, 8/12) and inhibited (33%, 4/12) by touch. For the touch-excited population of neurons, IT cells (N = 8) and PT cells (N = 18) displayed similar average post-touch response latencies (12.9 ms for PTs and 13.6 ms for ITs). However, ITs’ median response latencies (8.5 ms) were faster than PTs’ (11.5 ms) (**Figure 5C**), suggesting that both cell populations receive early input from thalamic or local thalamo-recipient cells (Pluta et al., 2015; Petreanu et al., 2009; Feldmeyer et al., 2005), though the difference between the two cell types remains non-significant (p = 0.48, Two-Sample Kolmogorov-Smirnov Test). The mean duration of the response was also similar between the two groups (PTs: 17.6 ± 6.36 ms C.I.; ITs: 15.1 ± 8.95 ms C.I., p = 0.87, Two-Sample Kolmogorov-Smirnov Test) (**Figure 5D**). PT cells displayed an average higher number of baseline-subtracted touch-evoked spikes than IT cells (PTs’ 0.15 ± 0.026 spikes/touch vs ITs’ 0.013 ± 0.009 spikes/touch) (**Figure 5E**). However, IT cells that were touch-excited displayed an overall higher percent-increase in spiking compared to PT cells (PTs: 101% mean increase above-baseline firing rate, ITs: 149% mean increase above-baseline firing rate; **Supplemental Figure 5A, B**), though the difference was not significant (p = 0.12, Two-Sample Kolmogorov Smirnov Test). However, this mean difference was enhanced when comparing only the first touches in trials (PTs: mean 341% vs.

**Figure 5.**
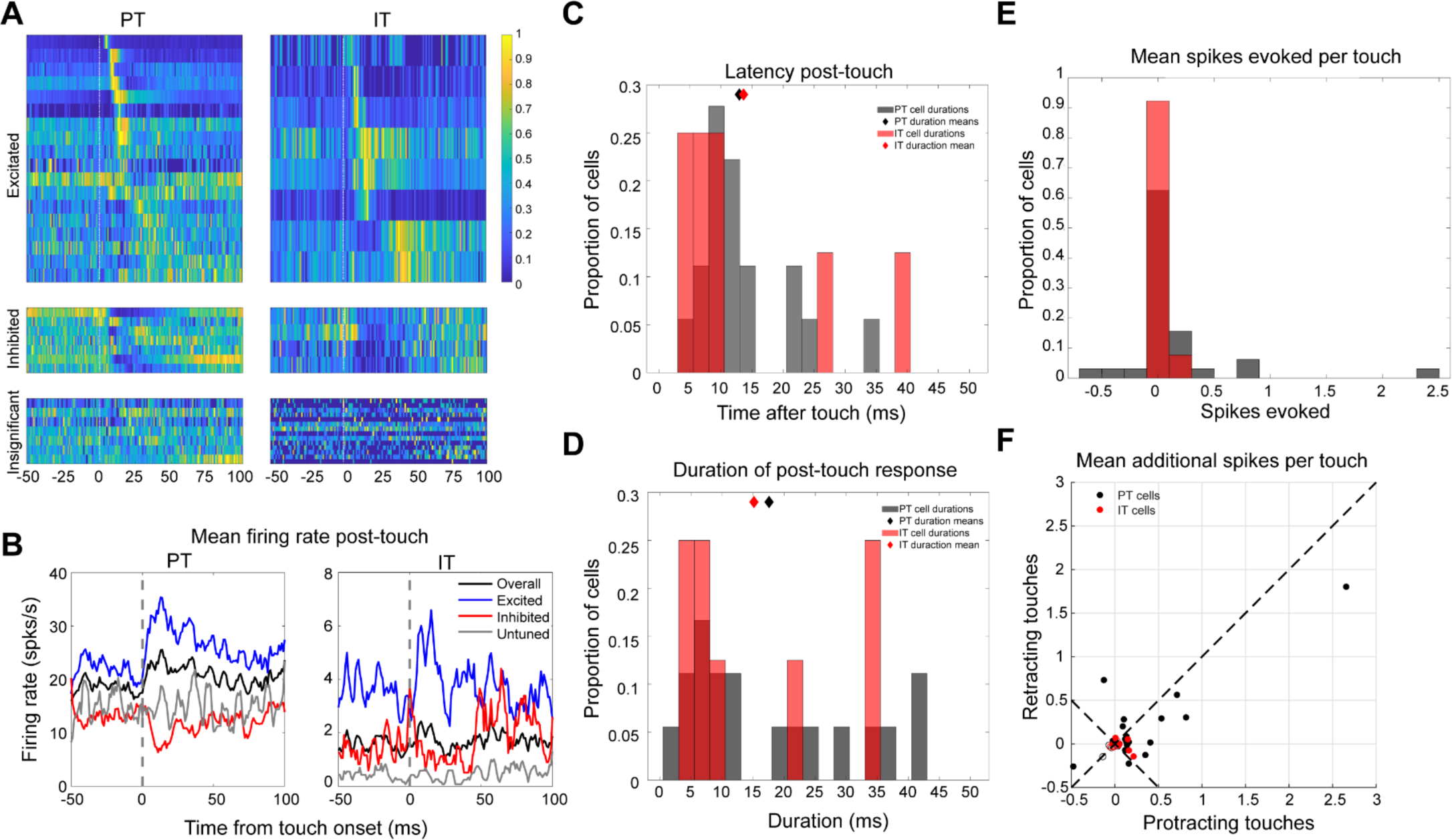
Touch-evoked Spiking Responses. (A) Heatmaps (PTs, left column; ITs, right column) of touch responses aligned to touch times. Cells that were considered ‘excited’ by touch were plotted on the top, cells inhibited by touch were plotted directly below excited cells, and any cells that had responses that were either not discernible or not significant (See Methods and Materials) were plotted at the bottom. (B) Population mean firing rate traces (PTs, left column; ITs, right column) for excited cells (blue), inhibited cells (red), untuned cells (gray), and all cells (black) aligned to the time of touch (gray dashed line). (C) Latencies for first spikes after touch for PTs (mean = 12.9 ms, black) and ITs (mean = 13.6 ms, red),p = 0.48, k-statistic = 0.33, Two-Sample Kolmogorov-Smirnov Test. Means are shown as diamonds in each cell type’s respective color. (D) Duration of the touch response is defined as the width of the response window. PTs are shown in black (17.6 ms ±6.26 ms C.I) and ITs in red (15.1 ± 8.95 ms C.I.), with means displayed as diamonds of each cell type’s color. A Two-Sample Kolmogorov-Smirnov Test was used to compare the two distributions: p = 0.87, k-statistic = 0.24. (E) Mean spikes evoked per touch for PTs (black, 0.15 ±0.026 spikes/touch, N = 32) and ITs (red, 0.013 ±0.009 spikes/touch, N = 26). (F) Mean additional spikes per touch for protracting versus retracting touches for PT cells (black, mean = 0.09 Δ spikes, p = 0.06,z-value = 1.87, signed rank = 232) and IT cells (red, mean = 0.06 Δ spikes p = 0.20, signed rank = 56).

IT mean 1361%), though again these two distributions were still not significantly different given the smaller sample size (p = 0.25, Two-Sample Kolmogorov Smirnov Test). Protraction touches elicited more spikes from both PT and IT populations compared to retraction touches, but not significantly so (PT: mean = 0.09 Δ spikes, p = 0.06, ITs: mean = 0.06 Δ spikes, p = 0.20, Wilcoxon Signed-Rank Test) (**Figure 5F**). Overall, PT and IT cells displayed similar spike latencies and post-touch response durations, though PTs produced significantly more above-baseline spikes compared to ITs for all touches, with ITs showing a consistently higher percent-increase in their firing rates for all touches but especially first touches. The prominence of the response to first touches in the IT population may allow touch events to stand out more strongly than in the PT population’s higher basal firing rate, making them more sensitive to rapid state changes in the sensory input or context.

### Adaptation to Touch

Strong adaptation is a known attribute of IT cells in brain slice (Hattox & Nelson, 2007; Guan et al., 2015) and in response to passive whisker deflections at naturalistic frequencies in rodents (Kheradpezhouh et al., 2017). We investigated if this attribute is maintained in the context of active touch behavior. We identified examples of adaptation in each cell type, with a significant drop-off in evoked spikes between first and subsequent touches (all touches after the first) in a trial (**Figure 6A, Supplemental Figure 6**). This was a general characteristic of most touch-excited cells in both populations (16/18 PT; 6/8 IT) (**Figure 6B, Supplemental Figure 7**). The mean adaptation ratios (mean subsequent touch evoked spikes / mean first touch evoked spikes) between the cell types were similar (0.62 for PTs, 0.57 for ITs, Two-sample Kolmogorov Smirnov Test) (**Supplemental Figure 8**). IT neurons adapted more sharply than PTs, as shown by the smaller evoked spike ratio between the first touch and second bin of subsequent touches (**Figure 6C**). However, this effect was not significant (p = 0.33, Two-sample Kolmogorov-Smirnov Test). Expressing adaptation in terms of inter-touch interval (ITI; the time since prior touch) showed a monotonically increasing response (**Figure 6A, D, E**). Overall, touch-excited PT and IT cells show strong adaptation to touch in the context of single-whisker tactile exploration, providing a relatively short window of time to gather information about the characteristics of the touched object.

**Figure 6.**
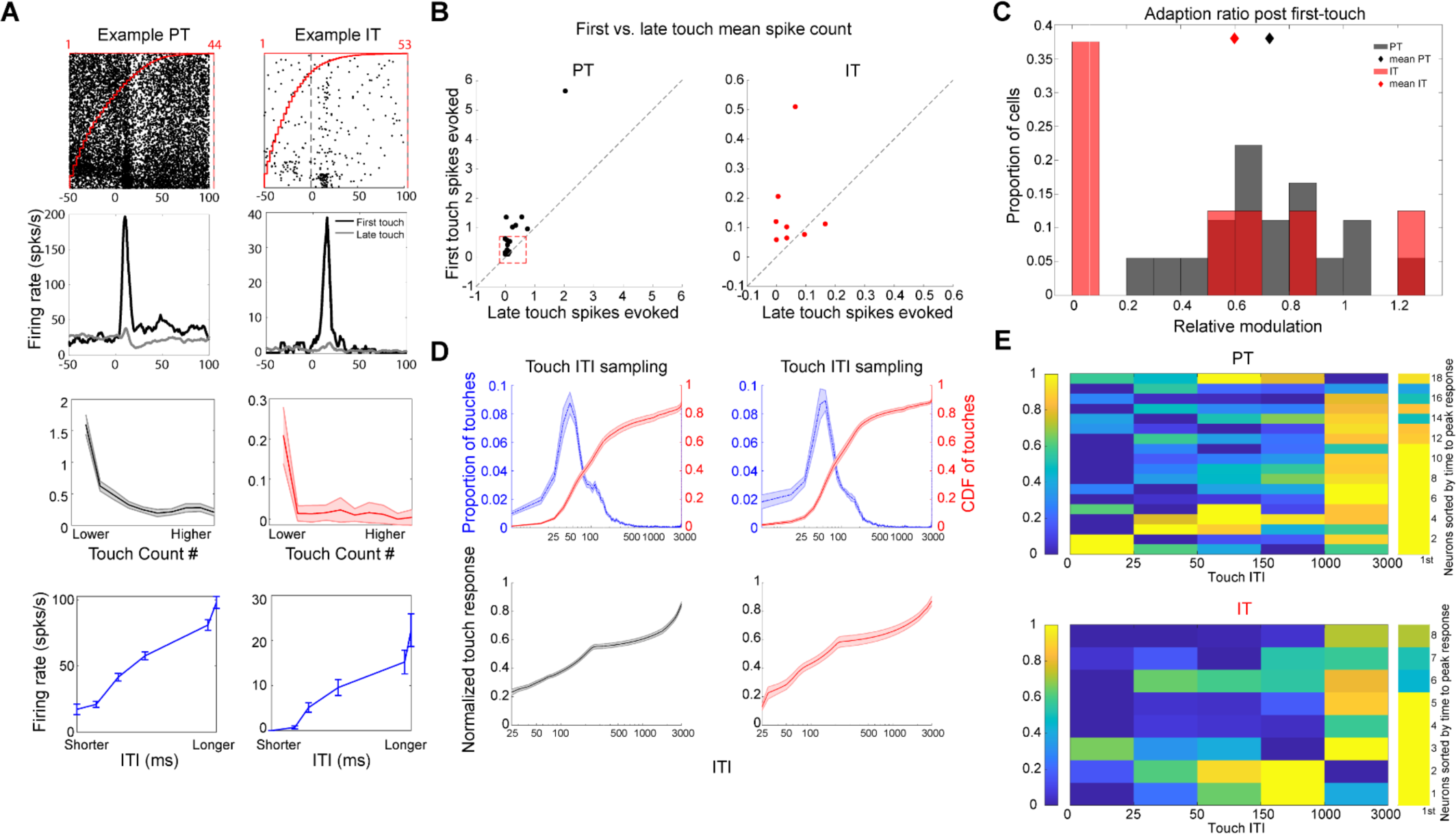
Adaptation to touch count and Inter-touch-interval. (A) Raster example (top) of a PT (left) and IT (right) sorted by touch order. (Upper-middle) PSTHs for the same example PT (left) and IT (right) as above. First-touch only is labeled in black, successive touches in gray. (Lower-middle) Adaptation of the example cells’ respective responses with successive touches from the first touch starting at the left of each figure and proceeding to the right. (Bottom) Each example cell’s responses to touch following different inter-touch-intervals. (B) First vs. late touch mean spike count. PTs are displayed on the left as black dots (mean first = 0.43 spikes/touch; mean late = 0.11 spikes/touch]). ITs are displayed on the right as red dots (mean first = 0.03 spikes/touch; mean late = 0.01 spikes/touch]). The red dashed box on the left panel indicates the space occupied by the ITs shown on the right panel. (C) Adaptation to touches immediately proceeding after first touch for PTs (black, 0.73) and ITs (red, 0.60), p = 0.33, k-statistic = 0.38, Two-Sample Kolmogorov-Smirnov Test. (D) (top-left) Proportion of touches found for PT cells (blue) at different ITIs, cumulative distribution of all touches across ITIs (red). (top-right) Same as in the top-left, but for IT cells. (bottom-left) Normalized mean touch responses for all PTs across different ITIs. (bottom-right) Same as in the bottom-left, but for ITs. (E) Heatmap of binned, normalized responses to touch ITI (top, PT, N = 18; bottom, IT, N = 8).

### Object Location Responses

Layer 5 neurons encode the azimuthal position of touched objects (Ranganathan et al., 2018; Cheung et al., 2020). We hypothesized that this encoding would be specific to PT neurons, based on their circuit connectivity. However, sorting touch evoked spike rasters by pole position revealed members of both populations that showed enhanced firing in particular regions of the azimuthal plane along the face (**Figure 7A**). Both PT (N = 32) and IT (N = 26) preferred locations tiled the space from posterior to anterior pole positions when considering all cells (**Figure 7B**). When comparing the two populations’ modulation widths, the IT population displayed a sharper response on average for the preferred pole location (**Figure 7C**).

**Figure 7.**
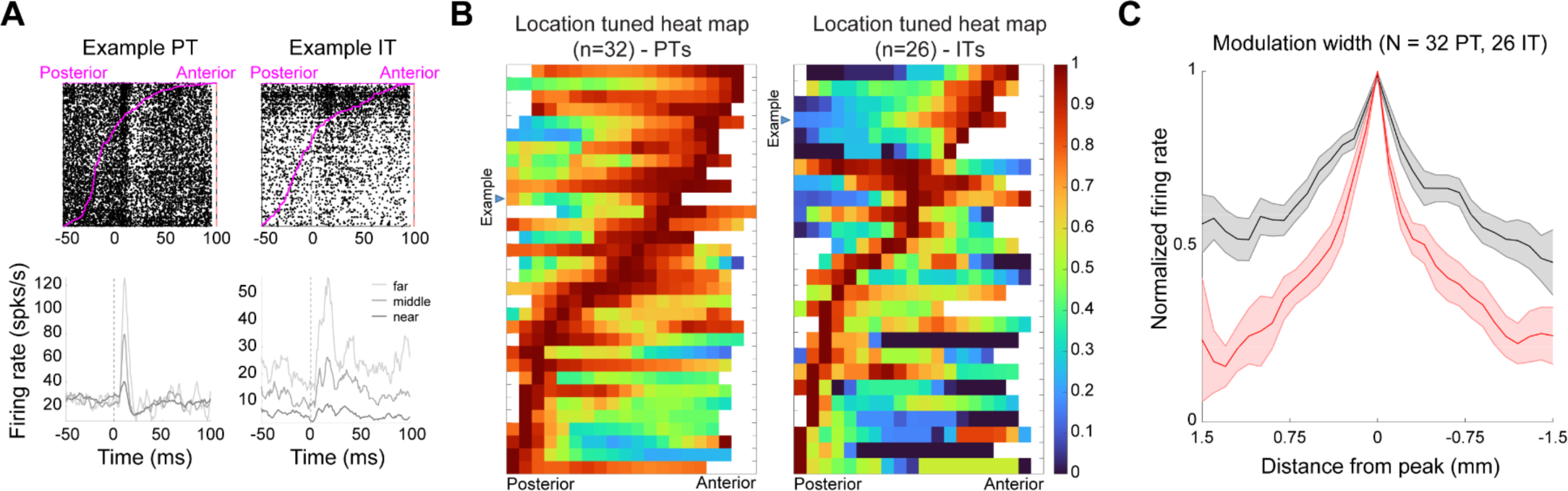
Cell-type Specific Responses to Object Location. (A) Touch-aligned rasters for an example PT (top-left) and an example IT (top-right) sorted by pole location from posterior to anterior (order shown in magenta). Touch-aligned PSTHs for each respective example (bottom), broken up by near (dark gray), middle (gray) and far (light gray) locations. (B) Population heat map of normalized object location responses for PTs (left) and ITs (right), each sorted by cells’ preferred locations from posterior to anterior. White spaces are insufficiently sampled pole locations. (C) Shape of normalized tuning curves across units from each cell type (PTs in black, ITs in red) indicating the mean peak response at preferred locations.

The accuracy of decoding information from population spikes is influenced by rates — higher rates provide more bit depth — and precision — sharpness of individual tuning curves. The PT population provides many more spikes per touch than IT, while the average IT neuron is more sharply tuned (**Figure 7C**). Which population can be used to extract the position of objects from their touch-evoked spike patterns more faithfully? Previously, the responses of 25 unlabeled L5 excitatory cells were adequate to predict a pole’s actual location across space at 60.5% accuracy at ≤ 0.5mm distance (Cheung et al., 2020). We applied a similar approach (multinomial generalized linear model; GLM) to each cell class. Models built with PT cells (**Figure 8A, D**) performed significantly better than their IT cell-only counterparts (**Figure 8B, D**) (PTs: 20.3 / 55.7 / 72.7% vs. ITs: 16.8 / 50.0 / 68.3% at ≤ 0 / 0.5 / 1mm from the actual location, 25 cells each). There is a possibility that ITs and PTs work in concert to construct a representation of object location in S1. Therefore, we also trained a model that drew from both datasets. The combined model performed slightly less well than either PTs or ITs alone (16.3 / 48.6 / 65.5% at ≤ 0 / 0.5 / 1mm from the actual location, 25 cells) (**Figure 8C, D**), indicating a lack of complementarity in their encoding. In summary, both cell types can be used to decode pole position with an accuracy ∼70% from 1 mm of actual position with 25 cells, and a pool of cells drawn from a combined population of PT and IT cells does not yield better performance at decoding object location. This suggests that they both receive adequate single-whisker motor and touch information to construct a representation of object location, but that they may do so in parallel and potentially for projection-specific purposes.

**Figure 8.**
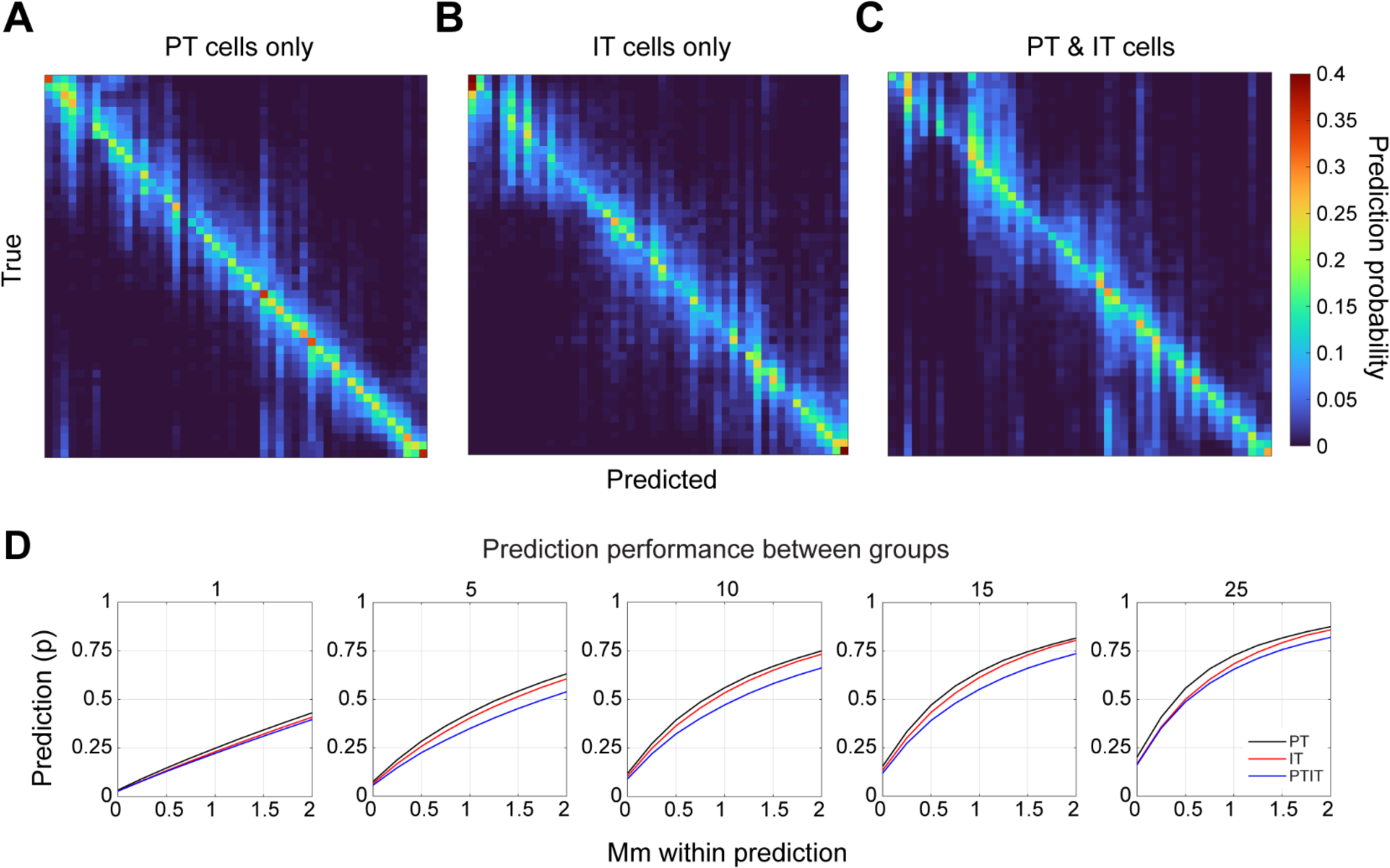
Decoding of Object Location by Cell Type. (A) Contingency tables for pole location decoding performance for PTs; true position of a pole is plot against the predicted position of a pole. N = 25 cell pool (B) Same as (A), but for ITs. (C) same as (A), but for a combined PT and IT set. (D) Successive subplots with varying cell counts used in the decoding step starting with 1 member in the first plot, and proceeding to 25 members in the last plot. PT-only decoding is plotted in black, IT-only in red, and mixed PT & IT in blue.

## Discussion

We used mouse lines with cell-specific expression in L5 of S1 to disentangle sensorimotor representations during a whisker-mediated task (**Figure 1**). We found that PT cells’ absolute modulation in response to active whisking was much higher, but ITs had higher relative modulation (**Figure 2**). The free-whisking kinematics of PTs and ITs were similar, with angle and amplitude modulation more correlated than phase and midpoint modulation across both cell subtypes (**Figures 3 & 4**). The touch-evoked absolute spiking of PTs was significantly higher than ITs (**Figure 5**). Both ITs and PTs adapted to multiple consecutive touches and touches following long ITIs elicited the largest touch-evoked spiking. However, several ITs showed higher adaptation between the first touch and touches immediately following (**Figure 6**). Both cell types displayed object-location-dependent firing rate changes upon touch, with PTs’ responses being broader but more accurate at decoding the actual location of the pole (**Figures 7 & 8**).

Our opto–tagging approach (Muñoz et al., 2014) allowed us to draw clear distinctions between two major output classes of S1 neurons; however, our methods had limitations. We did not recover individual cell morphologies (Egger et al., 2020) due to recording over multiple days. This knowledge could help in further assessment or sub-classification of the quickly adapting response seen in touch-responsive IT cells (Udvary et al., 2022; Oberlaender et al., 2012), and would also act as a robust secondary identifier of cell type beyond opto-tagging. Synchronous optogenetic activation of IT neurons could potentially recruit and misidentify PT neurons through the unidirectional projection from layer 5 ITs to layer 5 PTs (Kiritani et al., 2012; Brown & Hestrin, 2009; Anderson et al., 2010). We addressed this through the stringent co-application of a short latency threshold and a waveform-based criterion for inclusion (see Materials and Methods). However, this strict criteria may have excluded neurons with longer recruitment latency or idiosyncratic waveforms, which could produce bias in our assessment of cell-type population firing rates and other spiking characteristics. Finally, both of the transgenic lines used are corticostriatal (Gerfen et al., 2013, Morgenstern et al., 2022; Papale et al., 2023) and neither label cortico-cortical non-striatal excitatory IT neurons of L5, which may have distinct firing attributes (Rojas-Piloni et al., 2017; Kim et al., 2015).

Several of the individual findings in this dataset came as a surprise within the context of what was known of ITs’ and PTs’ respective roles within the S1 circuit. Many studies report ITs as relatively insensitive to whisker touch (Egger et al., 2020; de Kock et al., 2021) and preferentially responsive to whisking without touch (de Kock & Sakman, 2009). However, ITs were only relatively more modulated by whisking than PTs (**Figure 2**). In a few cells, they displayed robust touch responses relative to the baseline rate and adapted within one subsequent touch back to near baseline levels (**Figure 5, 6**). This later finding was peculiar for a couple of reasons. First, the timescale of the response was similar between ITs and PTs (**Figure 5C**), which indicates that ITs were receiving either early direct lemniscal thalamic input as in PTs (Bureau et al., 2006; Petreanu et al., 2009) or from direct combined excitatory (Feldmeyer et al., 2005) / inhibitory (Pluta et al., 2015) descending input from L4. The alternative of ITs receiving direct paralemniscal input (El-Boustani et al., 2020; Groh et al., 2014; Ohno et al., 2012) seems unlikely, as the timescale would be too long (Diamond et al., 1992; Sosnick et al., 2001) to appear with the low latency shown here. A potential explanation for these touch responses may come from rapid, brief excitation by layer 6 corticothalamic (CT) cells, which weakly disynaptically inhibits L4 and excites L5a regular-spiking (RS) and fast-spiking (FS) inhibitory cells (Kim et al., 2014). This initial net excitatory influence on L5a could shape a transient touch response in L5a ITs that gets abolished as descending, thalamically-driven L4 inputs inhibit them (Pluta et al., 2015). It is still unclear how these particular touch responses in ITs arise or what their specific function is.

The observation that actual object location was decoded as accurately as it was by a model trained with IT inputs also came as a surprise (**Figure 8B**). ITs performed more poorly than PTs at every distance from the actual pole position and at every size of neuron pool used in the decoding step, but generally only by ∼5-10% accuracy. Since the circuit already has PT neurons, which can seemingly do a similar computation more accurately, what could ITs be doing? ITs have been implicated in learning, forming internal models, and preparatory behavior (Salaj et al., 2021), and a recent study showed that learned-but not unlearned-motor behaviors were robustly represented in IT cells of M1 (Shinotsuka et al., 2023). It is possible that ITs’ sharp encoding of preferred object location is partially learned and changes through exposure to behaviorally relevant tasks and stimuli. With a more longitudinal recording method, this could likely be readily tested.

Overall, we conclude that PT neurons decode preferred object location more accurately and respond more robustly to touch stimuli than IT neurons despite displaying more adaptation than expected (**Figure 6C, E)** (Mason & Larkman, 1990). The responses of the PT cells in this dataset are largely consistent with a rate coding system, with more intense responses occurring closer to preferred pole positions and smoothly decreasing in magnitude with distance. ITs from this dataset, on the other hand, seem to be best suited for signaling fast contextual changes (*e.g.* onset of touch) due to the low number of touch-evoked spikes and lower background firing rate. Future work would benefit from adding multi-unit recording techniques that, in tandem with known behaviors of single units, could be leveraged to make more specific claims about the population-level behavior of each cell type, such as the potential use of synchrony-based location representations.

## Materials and Methods

### Lead contact and materials availability

Further information and requests for resources and reagents should be directed to and will be fulfilled by the lead contact, Samuel Andrew Hires.

### Ethics statement

All procedures were approved under USC IACUC protocol 20788 per United States national guidelines issued by the Office of Laboratory Animal Welfare of the National Institute of Health.

### Experimental model and subject details

Fourteen Sim1-cre x Ai32 crosses specifically expressing channelrhodopsin-2 (ChR2-H134R) in corticostriatal pyramidal tract neurons, and nine Tlx3-cre x Ai32 crosses specifically expressing ChR2(H134R) in corticostriatal intratelencephalic tract neurons were used in all experiments.

Individuals within the used population included males and females; all subjects were at least three months of age. A head-plate implantation procedure was conducted as described by Guo et al., 2014. Postoperatively, mice were housed with littermates where possible or singly housed if complications such as fighting occurred. Mice were provided food ad libitum and water restricted to 1 mL daily for one week before training and recording. A daily health assessment was completed to ensure that the mice were healthy.

### Method details

#### Object-localization task

Mice were trained in a whisker-based go/no-go object-localization task. Using a single whisker (C2), water-restricted mice were motivated to whisk and identify the location of a smooth vertical pole (0.6-mm diameter) 7–12 mm lateral from the whisker pad. The pole moved along the anteroposterior axis across 12 mm and was positioned using stepper linear actuators with 99-nm resolution, 25-μm accuracy, and <5-μm repeatability (Zaber NA11B30-T4). To avoid potential ultrasonic cues associated with stepper motor movement, the pole position was jittered 0–127 micro-steps (0–25 μm) on each trial. A pneumatic linear slider (Festo) was used to raise the pole vertically into touch reach for each trial. The Festo also provided a sound cue on the pole presentation onset.

Specific pole locations rewarded mice with water (2–8 μL), punished mice with a time-out (2 s) or had no effect based on the mouse’s decision to lick or withhold licking. In a go/no-go paradigm, four trial outcomes exist. In a minority of sessions in which the animals were trained, the close posterior 6 mm of pole locations (go) were rewarded with water rewards upon licking (hit) or had no effect if mice withheld licking (miss). The far anterior 6 mm of pole locations (no-go) were punished with time-out (false alarm) or had no effect if mice withheld licking (correct rejection). For the remaining sessions, rewards and punishment were given regardless of the pole location—go trials and no-go trials had overlapping pole locations.

### Behavior, videography, and electrophysiology

Animal behavior, videography, and electrophysiology were synchronized and captured during task performance using Wavesurfer (https://wavesurfer.janelia.org/). A single computer running BControl (MATLAB 2007b) was used to initiate each trial of the object-localization task and synchronize video and electrophysiology recordings via a second computer running Wavesurfer. Trial onset triggered high-speed video capture of whisker motion (1,000 fps) and electrophysiology recording of single-unit activity (MultiClamp 700b).

Whisker motion was captured from an overhead view and spanned 3 s, spanning pole onset to response window. Video frames were acquired using an Adimec N-5A100-Gm/CXP-6 camera and an Edmund Optics 0.18X 1⁄2” GoldTL Telecentric Lens (Model # 52–258) under 940-nm illumination on Streampix 7 software. Whisker shape and position were traced and tracked using Janelia Farm’s Whisker Tracker (https://www.janelia.org/open-science/whisk-whisker-tracking). A mask was traced around the edge of the fur to reduce tracking noise. The whisker angle is quantified at the intersection between the mask and the whisker. The whisker midpoint, instantaneous phase, and amplitude were decomposed from the bandpass- and zero phase– filtered (6–60 Hz, Butterworth) whisker-angle time series using the Hilbert Transform (MATLAB 2020a/2023b: Hilbert). Whisking amplitude and phase are defined as the magnitude and phase angle (radians) of the Hilbert Transform of the whisker-angle time series, respectively. A phase value 0 is the most protracted location of the whisk cycle, π and −π are the most retracted positions, and the sign of +/− defines retraction or protraction whisking directions. The whisking midpoint is the filtered (6–60 Hz) difference between the whisker-angle time series and bandpass-filtered signal. Whisker curvature is the amount of bending of the whisker measured 3–5 mm lateral from the whisker mask. The precise millisecond of touch was determined through an in-house auto-curation tool, Whisker Automatic Contact Classifier (WhACC) (Maire et al., 2023), which required a set of manually curated test images comprising whisker interactions with the pole and whisking in space to achieve high performance.

### In vivo loose-seal juxtacellular recordings

All animals used in this study were adult male or female transgenic mice (Sim1-Cre x Ai32 or Tlx3-cre x Ai32) expressing channelrhodopsin in either pyramidal tract (PT) cells or intratelencephalic (IT) cells. Following head-plate surgery, mice were trimmed to one whisker (C2), and intrinsic signal imaging was used to target the associated barrel column. A single whisker was maintained throughout training and recording. Prior to recording, animals were anesthetized (2% isofluorane), and a small craniotomy (200–300 μm) was made above the barrel column associated with the C2 whisker. On the first day of recording, animals were allowed to recover for one hour before recording. Recordings were repeated for 5.8 ± 5.4 sessions (mean ± SD) per animal.

To sample single-unit spiking activity in a manner unbiased by firing rate, blind juxtacellular loose-seal patch recordings were targeted to neurons using patch pipettes (Warner Instruments; 5–8 MΩ) filled with 0.9% saline (Growcells). Electrical recordings (n = 135 neurons) were acquired and amplified using MultiClamp 700b and Headstage CV-7B. The pipette axis was aligned parallel to the C2 barrel column at 35°. To perform an unbiased sampling of putatively PT neurons or IT neurons, we recorded from any isolated unit. An isolated unit was identified by an increase in resistance to 13–20 MΩ. Once a unit was isolated, it was opto-tagged at 10Hz frequency for 1.5 ms at 10ms width with bursts of 473 nm light at 9-16 milliwatts (beam width ∼200-μm diameter at skull, UltraLasers Model CST-L-473 nm– 50—OEM). The beam was focused onto the recording site from overhead and was used to test whether the recorded unit was either a PT neuron (for Sim1-Cre x Ai32) or an IT neuron (for Tlx3-Cre x Ai32). Stimulation was conducted for 5 trials in PTs and 1-2 trials in ITs. The ITs were stimulated for fewer trials to avoid an undesired behavioral response from the mouse characterized by general agitation and distress. After opto-tagging, an isolated unit was maintained for up to 300 trials, and all cells were kept for further analysis if they had at least 70% of all possible pole locations represented in the session.

Inclusion criteria for PT and IT neurons proceeded in a two-step process. Neurons whose latency to optogenetic stimulation was <=2.3ms were cleanly separable into two non-overlapping groups using a support vector machine with peak-to-trough ratio and duration as classifier variables. We used this boundary on all neurons with <=5.6ms latency to optogenetic stimulation to parse the responsive neurons into PT, IT, and unclassified cell types.

### Quantification and statistical analysis

#### Defining touch-response window

A smoothed (Bayesian adaptive regression splines (BARS); Wallstrom et al., 2008) response −50 ms to 50 ms around touch was used to evaluate the touch-response window. This window was generated two ways: one with all touches and one with the first touch only for cases in which first touches are not captured adequately. The touch-response window was defined as any time point from 5 to 50 ms post-touch in the smoothed response that exceeded baseline (−50 to 0 ms pre-touch) ± the 95% confidence interval. Then, the response windows were manually curated to best fit the probability density function of touches in cases where the initial approaches again failed to capture the window of the response. Any cells not assigned a touch response window due to non-response were assigned a response window that was the median of all response windows within the dataset.

### Tuning curves

A tuning curve is a response (firing rate) as a function of a stimulus (e.g., whisker position). For a single neuron, 5% of sampled touches or 5% of total whisking time points were used to define a point along the touch-or whisking-tuning curve. This method ensured 20 equally sampled bins of stimulus (e.g., whisker position) and response (firing rates) values. The response is defined as the firing rate within the touch-response window for touch tuning. For whisking tuning, the same response window as touch was used. If a neuron was not tuned to touch, the median touch-response window across all neurons was used to evaluate whisking tuning. The median touch-response window is 10–40 ms post-touch. The stimulus value is defined as the median of the stimulus in each sampled bin. Response values are defined as the mean of the responses in each sampled bin. Tuning curves were generated by smoothing the stimulus and response values using BARS. Neurons that had mean whisking responses less than 1 Hz were not evaluated.

We used a two-step process to define whether a neuron was significantly tuned to a specific location. We first performed a one-way ANOVA at an alpha level of 0.01 to identify if any angle/position’s firing rate at touch or during free-whisking significantly differed from another angle/position’s. If a neuron passed this first test, we moved on to the second step of the evaluation. In the second step, we shuffled touch/whisking responses 1,000 times and evaluated F-values from a one-way ANOVA. If the observed F-value was above the 95th percentile of the shuffled population distribution of F-values, we deemed the neuron as tuned. This second evaluation further ensures that the tuning we observed was not due to noise in neural responses. A neuron was considered significantly location-tuned if it passed both tests. Tuning preference is the location of the peak response of the tuning curve. To define the width of the tuning, a multiple comparison test using a Tukey-Kramer-type critical value was used to identify the first bins in both directions that were significantly different from the peak value. If no bins were significant, no modulation width was defined. Max and min responses were calculated from BARS-fitted tuning curves. The absolute modulation depth was calculated as the min response subtracted from the max response (max-min). The modulation depth was calculated as the min response subtracted from the max response, divided by the min response added to the max response ((max-min)/max+min).

A Shapiro-Wilk test was used at several steps to assess normality; in most cases, normality could not be determined, and a non-parametric test was used to test significance.

### Neural decoding

We used multinomial logistic regression to decode pole location implemented using glmnet [61], as described in Cheung et al. 2020. Only touch units that sampled at least 70% of the pole location range were used for decoding. Each unit had a tuning curve that was interpolated to 40 bins to estimate location to 0.25-mm resolution. At each bin, 50 samples were drawn from a Poisson pdf with a λ as the mean of each interpolated bin. We justified drawing from a Poisson pdf because we found that at touch, the number of spikes generated in the touch-response window followed a Fano factor of 1.01 ± 0.29 for PTs and 1.03 ± 0.09 for ITs (mean ± SD, S3E Fig). For the design matrix, each row is a location bin, each column is a single neuron, and each entry is a sampled neural response for the associated neuron.

The decoder was run for 10 iterations. During each iteration, a random 70% of trials were allocated for training and the remaining 30% for testing. Lasso regularization (alpha parameter 0.95) was used to reduce overfitting. To identify the number of units required, we sampled varying numbers of neurons with replacement from the units used to train the original model 1000 times. The indices of the selected neurons were used to create a new population design matrix and matrix of learned coefficients from the original values. The prediction probabilities of location were computed using the same method as seen in Cheung et al. 2020. The predicted location was chosen as the location with the highest probability. Model evaluation of accuracy and resolution was performed on the test set. Model accuracy was defined as the total number of correct predictions divided by the total number of predictions. A confusion matrix made from true and predicted locations was normalized across the total number of given true cases and used to define the decoding resolution and neurometric curves. Decoding resolution is defined as the total number of predictions within n bins of the diagonal, where each bin was 0.25 mm.

---

**Supplemental Figure 1.**
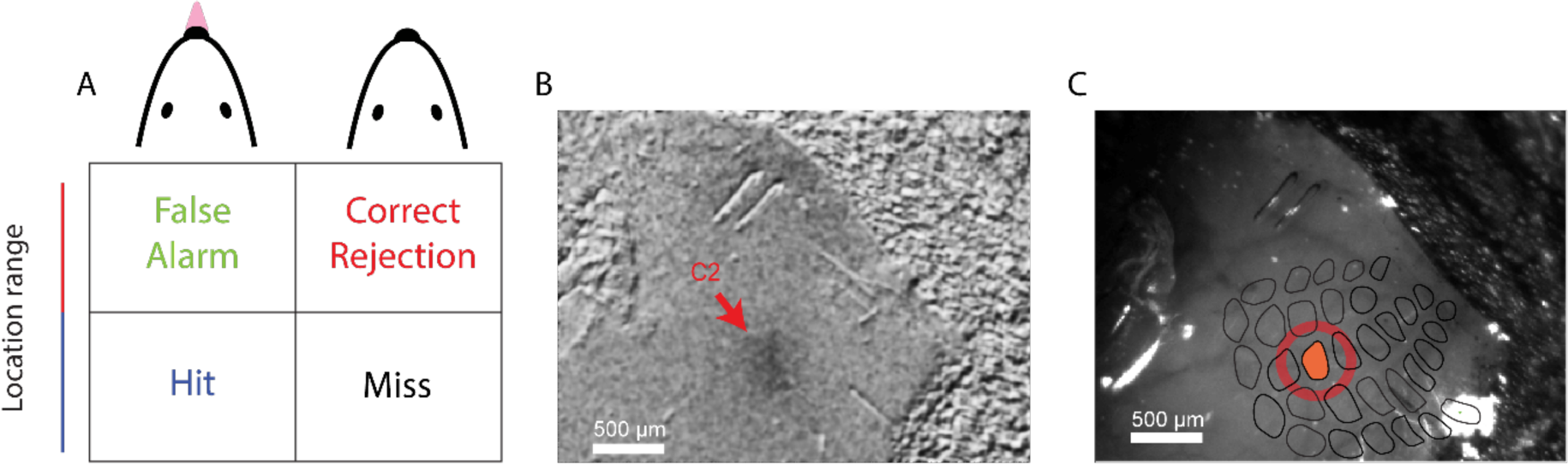
Intrinsic signal imaging. (A) Intrinsic signal imaging showing a highlighted region of activity during piezo-driven whisker stimulation. (B) Skull vasculature with a diagram of the barrel field overlaid and C2 highlighted in red (bottom).

**Supplemental Figure 2.**
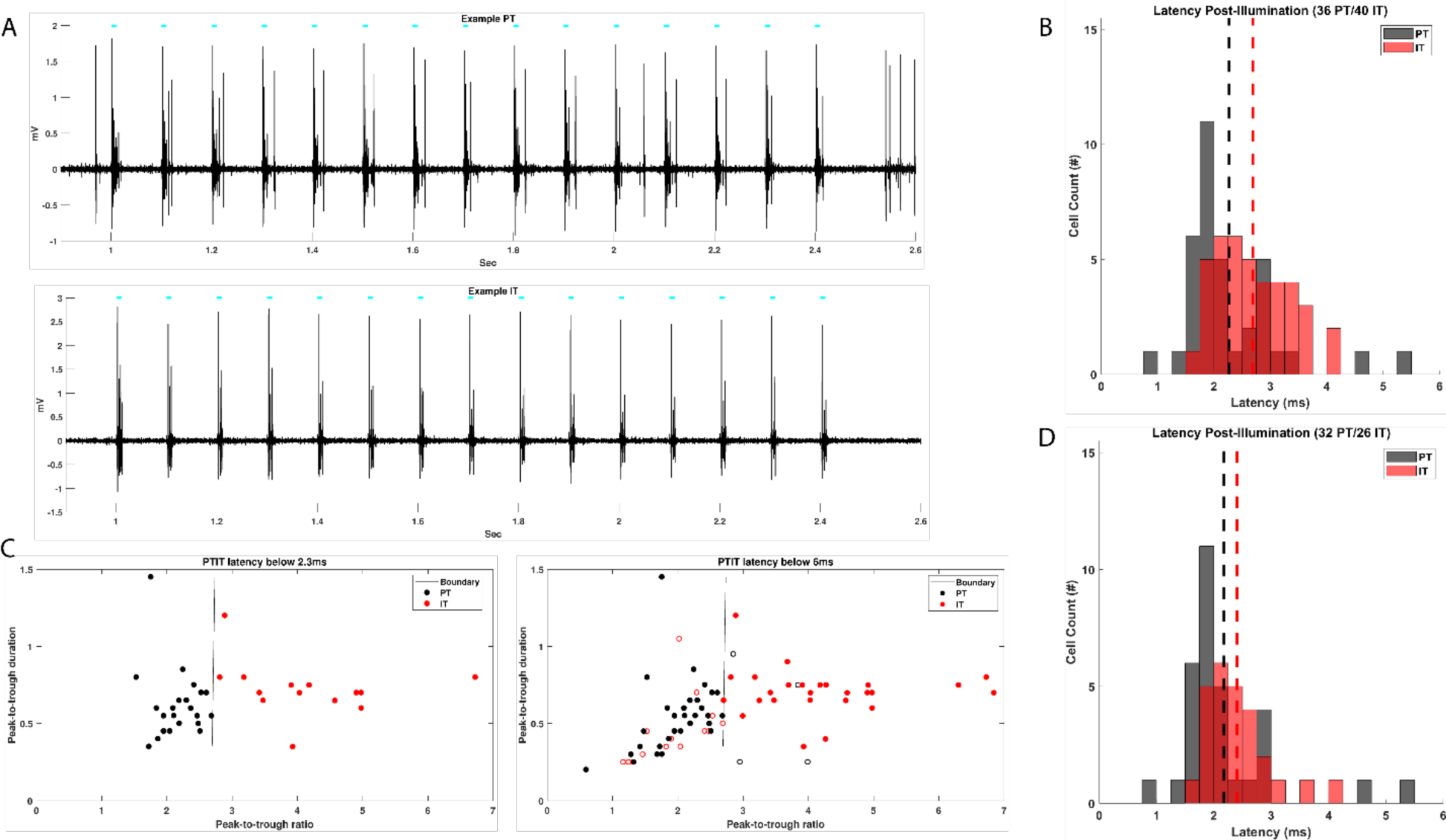
Opto-tagging results and waveform criteria. (A) Opto-tagging examples for PT (top) and IT (bottom) cells. Pulse times for 472 nm light are indicated by the cyan ticks above each raw trace. Light was pulsed at 10 Hz frequency at 10ms pulse width for 1.5s at 9-16 mW. (B) Initial Latency histogram for all cells retained after an initial 5.6ms time-to-first spike post illumination cut-off was imposed. PTs (N = 36) in black and ITs (N = 40) in red. Means are displayed as dashed lines of their respective colors.(C) Support Vector Machine (SVM) - based boundary for further classification of PT and IT cells by their spike waveform peak-to-trough ratios plotted against their peak-to-trough durations (left). (right) Cells retained in the population are shown as dots of their respective cell type colors (PTs, black, ITs, red). Open circles indicate cells for whom we were not confident defining as either one cell type or the other. (D) Latency histogram for all cells retained in the dataset with PTs (N = 32) in black and ITs (N = 26) in red. Means are displayed as dashed lines of their respective colors (PTs: 2.18 ± 0.87 ms, ITs: 2.41 ± 0.57 ms; mean+sd).

**Supplemental Figure 3.**
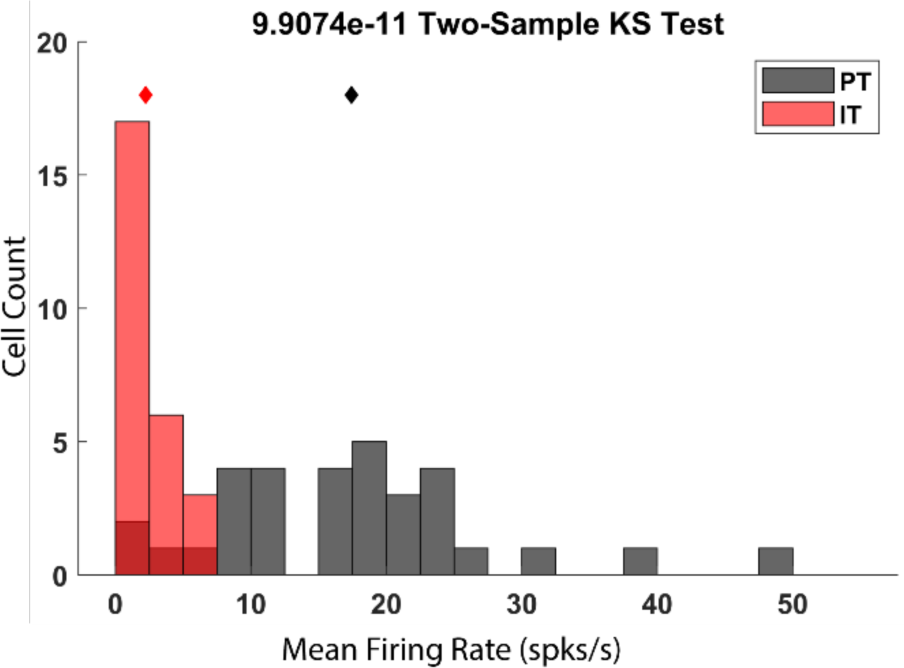
Session Mean Firing Rates. Population mean firing rate for PTs (black, 17.42 ± 3.53 spk/s, mean ± sd) and ITs (red, 2.23 ± 0.73 spks/s, mean ± sd). The Distribution of the two populations’ responses was tested with a Two Sample Kolmogorov-Smirnov Test: p = 9.91e-11, k-statistic = 0.88.

**Supplemental Figure 4.**
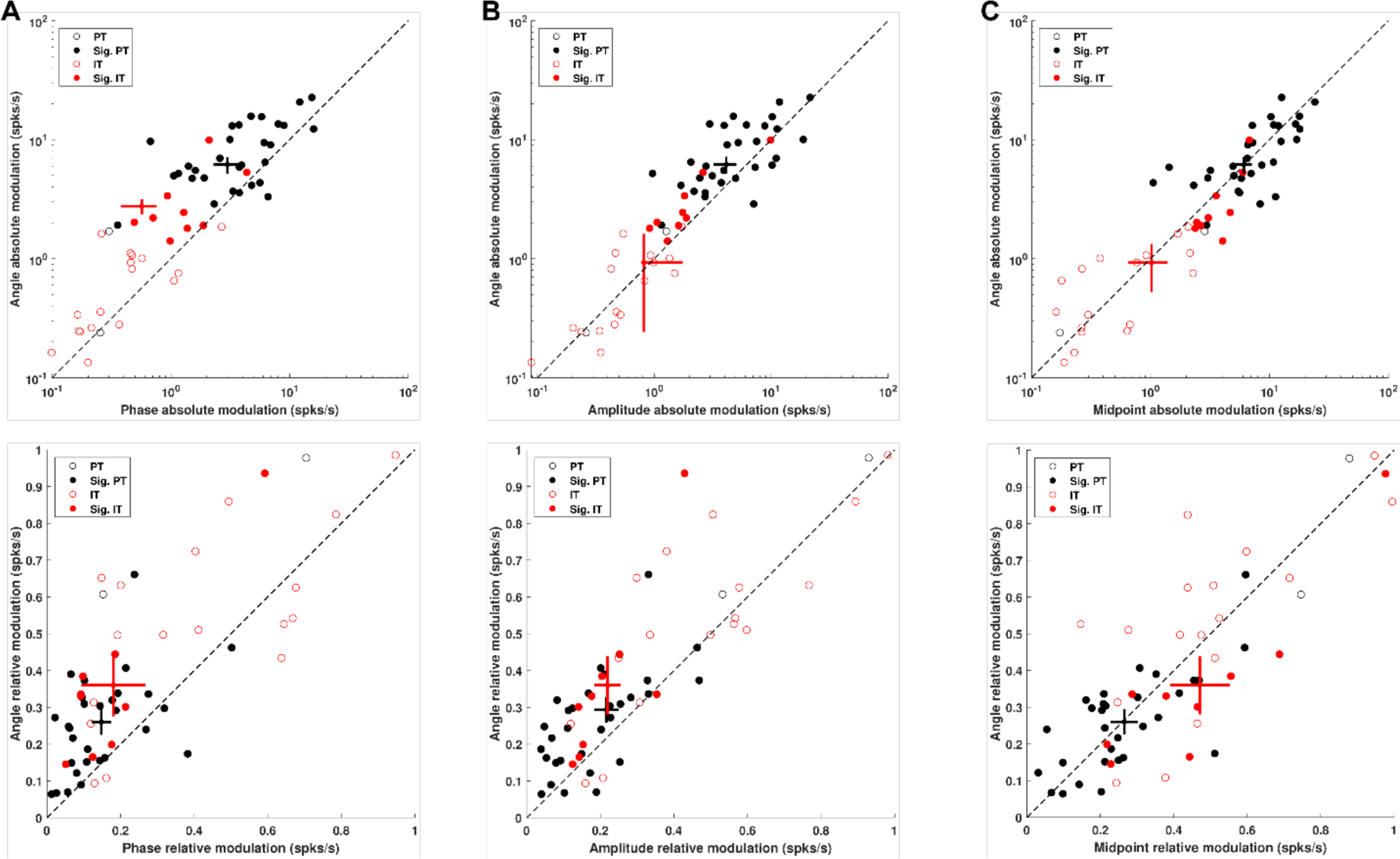
Absolute and Relative Angle Modulation vs Phase, Amplitude, and Midpoint. PTs are in black, and ITs in red, with significantly modulated units filled. All statistical comparisons were done using a Wilcoxon Signed-Rank Test. (A) (top) Angle absolute modulation plotted against phase absolute modulation, log scale (top). PT: angle geometric mean = 7.19 spks/s, phase geometric mean = 3.50 spks/s, p = 5.37e-05, z-value = 4.04, signed rank = 480. IT: angle geometric mean = 2.76 spks/s, phase geometric mean = 0.57 spks/s, p = 1.9e-03, z-value = 3.11, signed rank = 298. (bottom) Angle relative modulation plotted against phase relative modulation. PT: angle mean modulation = 0.29, phase mean modulation = 0.17, p = 3.59e-05, z-value = 4.13, signed rank = 485. IT: angle mean modulation = 0.47, phase mean modulation = 0.33, p = 2e-03, z-value = 3.10, signed rank = 297. (B) (top) Angle absolute modulation plotted against amplitude absolute modulation, log scale (top). PT: angle geometric mean = 7.19 spks/s, amplitude geometric mean = 4.71 spks/s, p = 1.9e-03, z-value = 3.10, signed rank = 430. IT: angle geometric mean = 2.76 spks/s, amplitude geometric mean = 1.89 spks/s, p = 0.08, z-value = 1.74, signed rank = 244. (bottom) Angle relative modulation plotted against amplitude relative modulation. PT: angle mean modulation = 0.29, amplitude mean modulation = 0.22, p = 4.71e-04, z-value = 3.50, signed rank = 451. IT: angle mean modulation = 0.47, amplitude mean modulation = 0.38, p = 0.02, z-value = 2.35, signed rank = 268. (C) (top) Angle absolute modulation plotted against midpoint absolute modulation, log scale (top). PT: angle geometric mean = 7.19 spks/s, midpoint geometric mean = 7.04pks/s, p = 0.64, z-value = −0.47, signed rank = 239. IT: angle geometric mean = 2.76 spks/s, midpoint geometric mean =3.69 spks/s, p = 0.07, z-value = −1.84, signed rank = 103. (bottom) Angle relative modulation plotted against midpoint relative modulation. PT: angle mean modulation = 0.29, midpoint mean modulation = 0.27, p = 0.99, z-value = −0.02, signed rank = 263. IT: angle mean modulation = 0.47, midpoint mean modulation = 0.47, p = 0.56, z-value = −0.57, signed rank = 153.

**Supplemental Figure 5.**
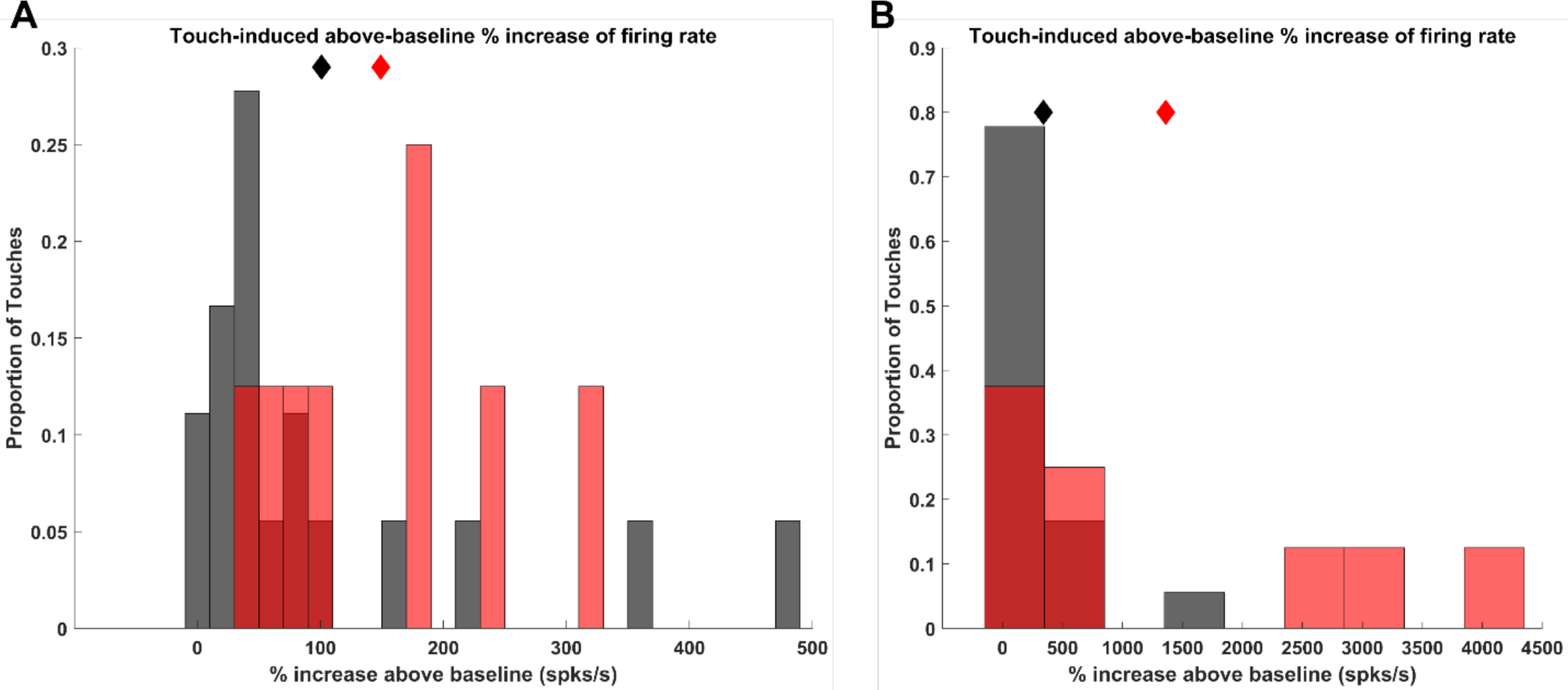
Percent-increase of firing rate above baseline. (A) Touch-induced above-baseline % increase of firing rate with PTs (mean increase of 101%) in black and ITs (mean increase of 149%) in red. Means are displayed as diamonds of each cell type’s respective representative color. Two-Sample Kolmogorov-Smirnov Test, test statistic = 0.47, p = 0.12. (B) First-touch-induced above-baseline % increase of firing rate with PTs (mean increase of 341%) in black and ITs (mean increase of 1361%) in red. Means are displayed as diamonds of each cell type’s respective representative color. Two-Sample Kolmogorov-Smirnov Test, test statistic = 0.40, p = 0.25.

**Supplemental Figure 6.**
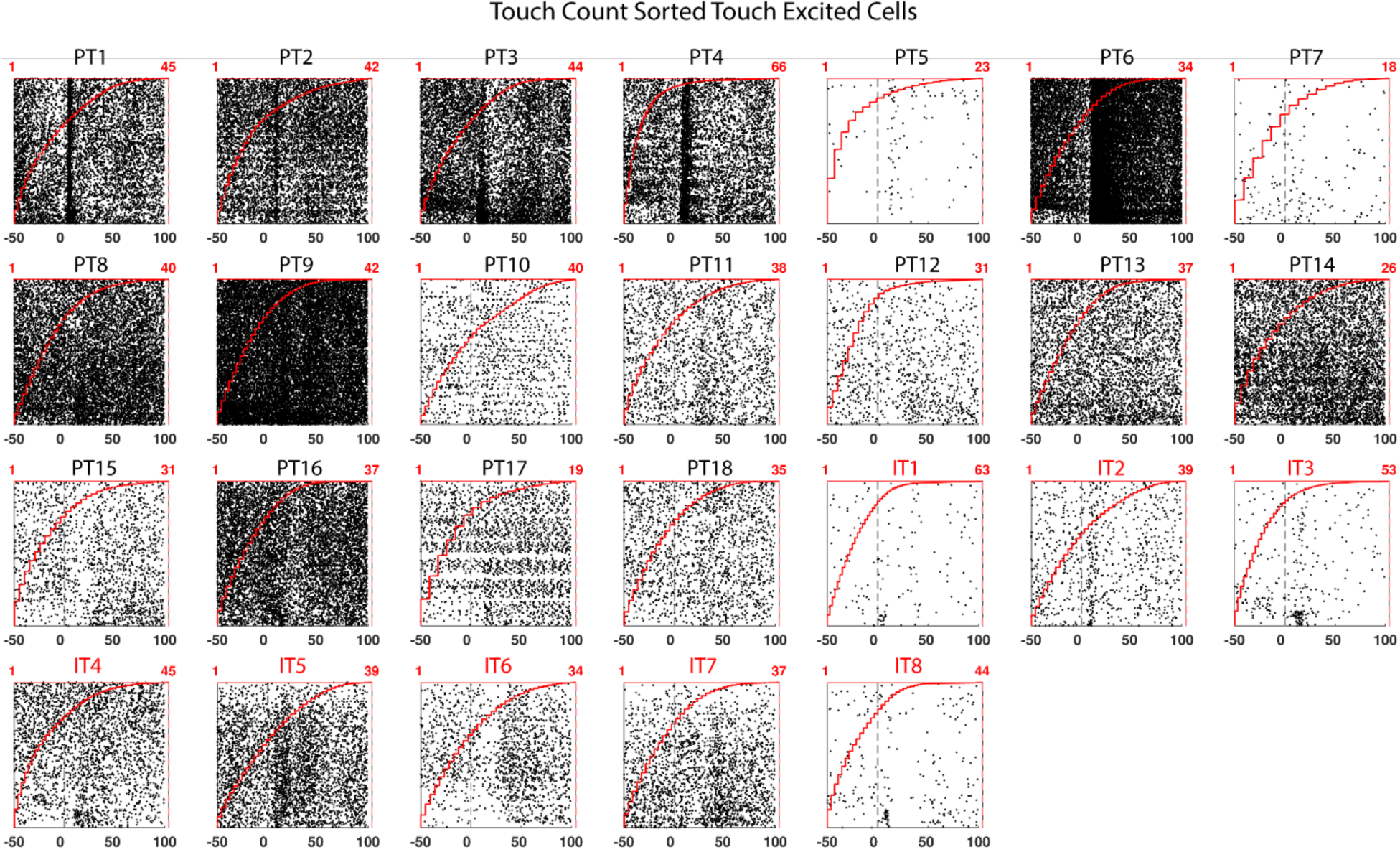
Touch-count-sorted rasters for significantly touch-responsive cells. Touch-count-sorted and touch-aligned spike rasters for PT and IT cells, with first touches starting at the bottom of each raster. The red curve indicates which touch in a trial the raster is derived from. PT cell labels are shown in black, IT cell labels are shown in red.

**Supplemental Figure 7.**
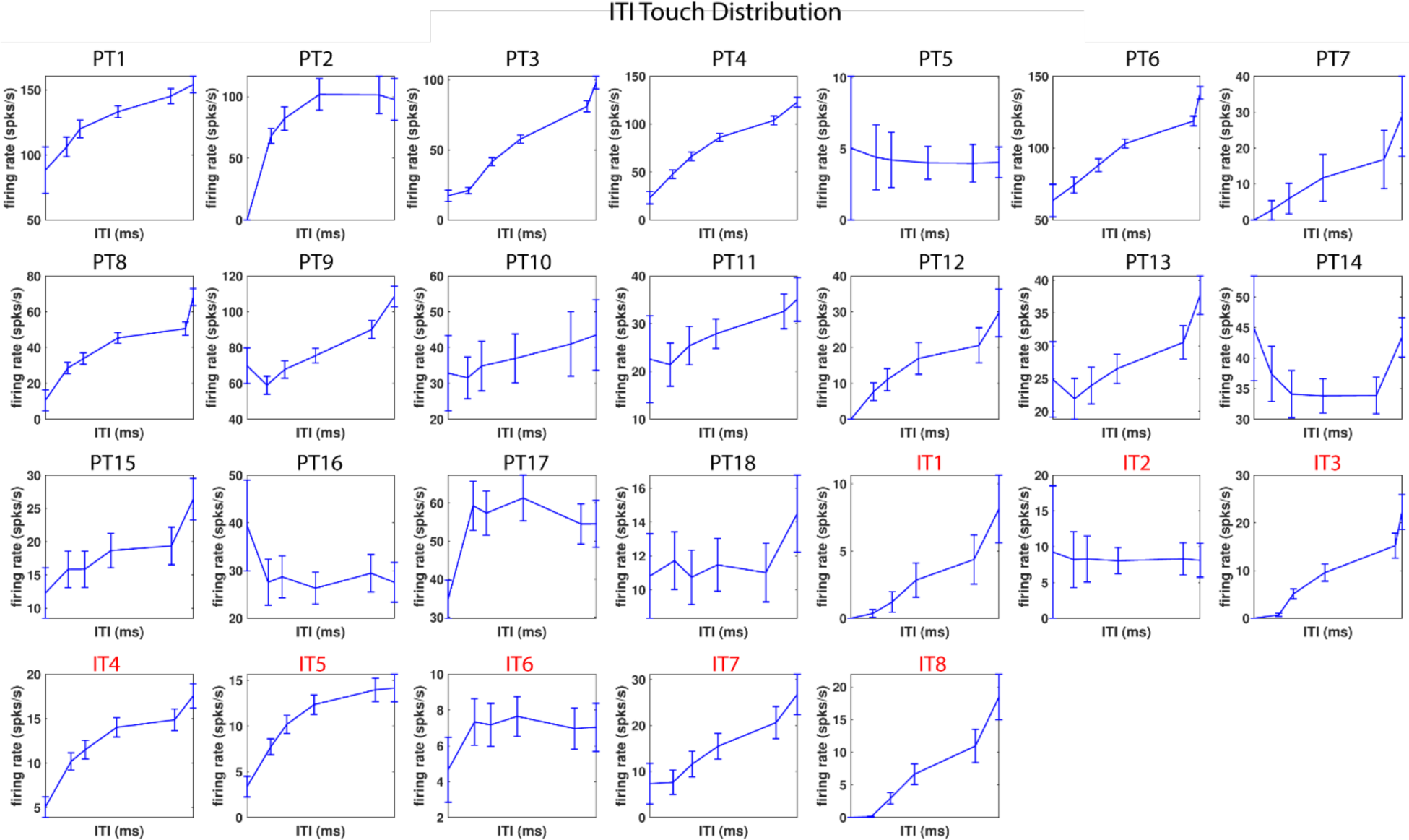
ITI touch responses for significantly touch-excited cells. Touch responses plotted against preceding ITIs for PTs (black cell label) and ITs (red cell label). The x-axis of each subplot starts at the left from the minimum observed touch ITI for that session and proceeds to the maximum ITI on the right.

**Supplemental Figure 8.**
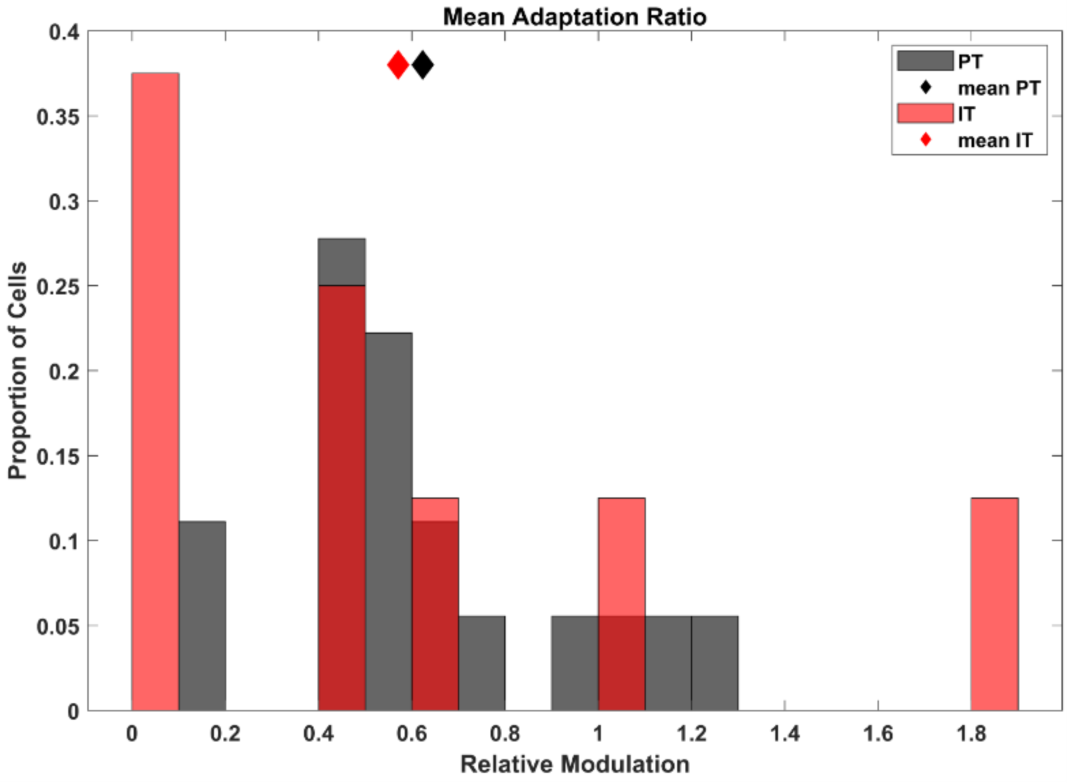
Mean adaptation ratios across all touch counts. Mean adaptation ratios for each cell type (PTs in black, 0.67, ITs in red, 0.57, p = 0.33, k-statistic = 0.38, Two-Sample Kolmogorov-Smirnov Test).

